# A Bayesian approach for identifying similar transcript dynamics using curve registration

**DOI:** 10.64898/2026.04.26.720911

**Authors:** Ruth Kristianingsih, Alexander Calderwood, Gurpinder Singh Sidhu, Shannon Woodhouse, Hugh Woolfenden, Smita Kurup, Rachel Wells, Richard J. Morris

## Abstract

Changes in gene expression over time can provide valuable insights into developmental processes and responses to the environment. Differences in expression may be indicative of potential differences in regulation. Comparing transcript dynamics may help identify correspondences between developmental stages within and between species, differences in the timing of key events during development, and transcriptional response to treatments or perturbations. A straightforward comparison between the dynamics is, however, hindered by measurements that were taken at different time points and over different timescales. To address this, we developed a statistical approach that seeks the optimal alignment between two time series as a function of a temporal shift and stretch. We validated our approach using simulated data and applied it to several transcriptome datasets, including comparisons between different plant species. Our development facilitates knowledge transfer from model systems to less studied species, the identification of modules of co-regulated genes, and the discovery of condition-specific, temporally differentially-expressed genes. The method is provided freely available as an R package.

## 1 Introduction

Analysis of gene expression over time can shed light on the inner workings of cells and provide insights into transcriptional regulation and gene function. Variation in gene expression can lead to phenotypic differences within and between species. A deeper understanding of this variation is, therefore, central to elucidate the mechanistic basis of differences in traits [1]. Changes in expression can be used to identify genes that might be associated with a specific environmental stimulus or developmental process, and to construct or validate gene regulatory networks [2–4]. The biological processes that lead to changes in expression may occur at different developmental times and at different rates in different species, strains, individuals, tissues, or conditions. Comparison of gene expression dynamics can help identify correspondences between developmental stages, differences in the timing of key events during development [5, 6], and transcriptional responses to biological, chemical or physical perturbations [7].

Comparing two different datasets requires a metric for quantifying how similar they are. Several such measures [8–10], rely on the datasets having corresponding time points, allowing the difference, often the Euclidean distance, between measurements at each time point to be computed. However, experimental sampling times between datasets often do not correspond, hindering such a straightforward comparison.

Several methods have been suggested to overcome these issues. One commonly used approach is dynamic time warping (DTW) [3, 5, 11]. DTW was first developed for speech recognition [12], but has been applied in many different fields, including bioinformatics, medicine, and engineering [12, 13]. This method establishes correspondences between different time points, *t*_*i*_ (1 ≤ *i* ≤ *N*_1_) and *t*_*j*_ (1 ≤ *j* ≤ *N*_2_), of two datasets (*t*_*i*_, *y*_*i*_) and (*t*_*j*_, *z*_*j*_) of size *N*_1_ and *N*_2_, by minimising the distance between the *y*_*i*_ and *z*_*j*_ values. The absolute distance, *d*(*y*_*i*_, *z*_*j*_) = |*y*_*i*_ − *z*_*j*_ |, is commonly used here. The smaller the summed distance over all corresponding time points, the more similar the dynamics. However, DTW does not answer the question of whether two time series can be considered the same, which is often of interest when comparing the expression dynamics of two genes. DTW matches the start and end indices of the first dataset with the corresponding indices of the other dataset, which can be problematic for pairs of gene expression profiles that have dissimilar patterns and exhibit differential progression over time. DTW matches every time point from one dataset to one time point from the other dataset, and *vice versa*. However, multiple points in one set can map to the same points in the other dataset, i.e. the correspondence need not be one-to-one, making the resulting alignment challenging to interpret biologically [14]. DTW4Omics [15] is based on DTW and uses a permutation test to estimate the significance of the alignment between two time series, but can be computationally costly for exploring all associations [16]. A recently developed R package, TimeMeter [13], performs post-processing of DTW results to quantify the temporal similarity in gene expression. However, the interpretability of the results remains challenging. Furthermore, these methods do not handle unbalanced sampling well or missing data points.

Several alternative approaches based on hidden Markov models (HMM) have been proposed [3, 14, 17]. HMM-based methods facilitate effective translation of the timeseries relative to one another but struggle when comparing datasets over different timescales. Additionally, these approaches can only detect positively correlated dynamics. On the other hand, DynOmics [16], based on the fast Fourier transform, can identify time shifts as well as positive and negative correlations between two profiles. The computational time of these approaches increases significantly when applied to large datasets.

Here, we describe an approach for aligning a query dataset (here, gene expression over time) to a reference dataset through the application of a suitable transformation, so that the transformed data maximally coincides with the reference data [18, 19]. This approach uses two parameters, stretch and shift factors, to explain differences between expression dynamics. A stretch factor can be interpreted as an increase or decrease in the rate of a biological process [5]. A shift factor represents the change in the relative timing, e.g. the delay, of a transcriptional response. In developmental biology, both the stretch and shift factors are known as heterochronic parameters, which describe a change in the relative timing of a process or a change in the rate [20]. The Bayesian Information Criterion [21, 22] is used to evaluate whether one model or two models best describe the datasets. One model suggests that the datasets are similar, two models that they are not. We tested our approach on simulated data and experimental expression time series. We show that our software can identify pairs of genes with similar expression dynamics, thereby inferring stretch and shift factors. These parameter values can be used to inform further analysis, such as distinguishing between homologous/paralogous genes that may have acquired new functions, identifying the timing of responses, and providing clues for the role of genes of unknown function. Whilst the method was developed for analysing gene expression profiles, there is nothing specific to this application in the approach, making it generally applicable to any data consisting of ordered values, such as time series.

## 2 Methods and Implementation

### 2.1 Two time series can be compared by evaluating how well they can be explained by a joint model

Suppose we have two sets of values at specified, but potentially different, time points, Figure 1. The different datasets could, for example, be expression values over time for the same gene under different environmental conditions or homologues in different species. We wish to compare these time series.

**Figure 1:**
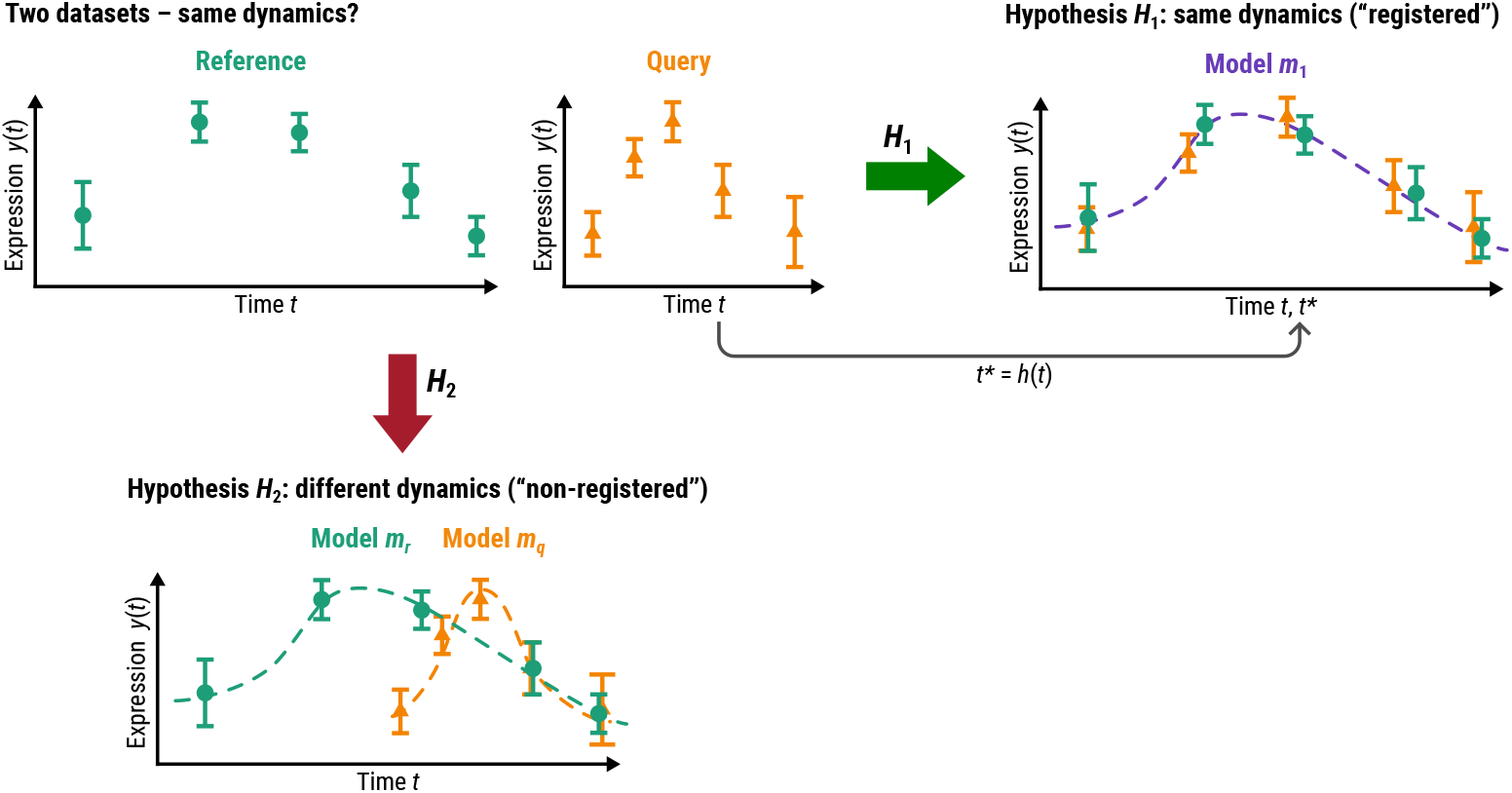
Evaluating whether two datasets with different time points exhibit similar dynamics depends on the choice of model. We depict two datasets: a reference against and a query dataset. The green (reference) and orange (query) expression values have different time points associated with them, hindering a simple comparison of their corresponding expression values at equivalent time points. We seek a transformation of time, *t*^∗^ = *h*(*t*), that makes the datasets as similar as possible. We can then ask whether the datasets are best explained by a single model (a function *m*_1_), or whether two models (functions *m*_*r*_ and *m*_*q*_) provide a statistically better explanation. We denote the situation in which one model best explains the data as Hypothesis *H*_1_ with Hypothesis *H*_2_ being the alternative hypothesis where two models are required to explain the data. The notation is explained in more detail in the Methods section.

We will refer to one time series as the *reference* (*r*) and the other as the *query* (*q*). We denote the time points of the reference (r) dataset by *t*_*r,i*_ and the corresponding values by *y*_*r,i*_, where 1 ≤ *i* ≤ *N*_*r*_ with *N*_*r*_ being the total number of time points in the reference dataset. The time points for the query (q) are denoted by *t*_*q,j*_, where 1 ≤ *j* ≤ *N*_*q*_ with *N*_*q*_ being the total number of time points in the query dataset. The measurements of the query dataset are denoted by *y*_*q,j*_. If multiple replicates are available then *y*_*r,i*_ and *y*_*q,j*_ can be used to capture the mean values at each time point and additional data descriptors, *σ*_*r,i*_ and *σ*_*q,j*_, to capture the estimated standard deviations. The time points can differ between query and reference.

If the time points *t*_*r,i*_ and *t*_*q,j*_ of two datasets are not the same, then the associated values *y*_*r,i*_ and *y*_*q,j*_ cannot be directly compared. In this case, an underlying model of the data is required. This choice of model impacts all subsequent inferences (see Supplementary Figure S1 and Supplementary Table S2). We denote the model for the reference dataset by the function *m*_*r*_(*t, θ*_*r*_) and the model for the query dataset by the function *m*_*q*_(*t, θ*_*q*_), where *θ*_*r*_ and *θ*_*q*_ represent the parameters in the models. It may be that one single model, a function *m*_1_(*t, θ*_1_) with parameters *θ*_1_, can explain both datasets simultaneously.

Determining whether two time series are within experimental error (i.e. one model can explain the data), or not (i.e. two models are needed to explain the data), can be cast in a probabilistic framework. The task then becomes to estimate the evidence for the two hypotheses:

- Hypothesis *H*_1_: the datasets are best explained by *one* common model *m*_1_(*t, θ*_1_)
- Hypothesis *H*_2_: the datasets are best explained by *two* different models *m*_*r*_(*t, θ*_*r*_) and *m*_*q*_(*t, θ*_*q*_).

To compute the probability of each hypothesis given the data we can employ Bayes’ theorem,

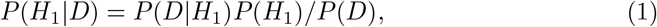

where *H*_1_ is the joint model for the datasets *r* and *q, P* (*H*_1_) is the prior probability for hypothesis *H*_1_, *P* (*D*|*H*_1_) is the marginal likelihood or evidence, and *P* (*D*) is the probability of the data (normalising factor). The model of *H*_1_ may have a number, 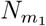 of parameters, *θ*. The evidence can be computed by integrating the likelihood, *P* (*D*|*H*_1_, *θ*_1_) over the prior of the parameters, *P* (*θ*_1_|*H*_1_), *P* (*D*|*H*_1_) = ∫ *P* (*D* | *H*_1_, *θ*_1_)*P* (*θ*_1_ | *H*_1_)*dθ*_1_. We assume Gaussian noise at each data point (*r, i* for the reference set, *q, j* for the query), leading to the following likelihood,

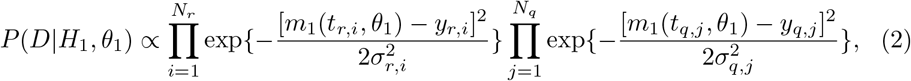

where *σ*_*r,i*_ and *σ*_*q,j*_ are the estimated standard deviations for the data points of the reference and query datasets at each time point *i*. The likelihood of *H*_1_, *P* (*D*|*H*_1_, *θ*_1_), depends on the parameters *θ*_1_ of the function *m*_1_(*t, θ*_1_) that is used to describe the reference and query datasets simultaneously.

For hypothesis *H*_2_ the likelihood depends on the parameters *θ*_*r*_ and *θ*_*q*_ of the models, *m*_*r*_ and *m*_*q*_, for the different datasets,

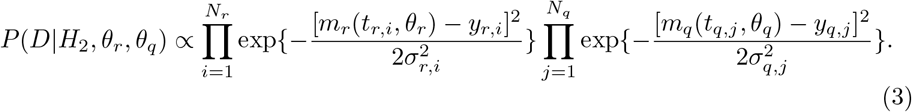

With the above likelihood functions, we can compute the marginal likelihoods by integrating over the parameters, *θ*_1_ for *H*_1_ and *θ*_*r*_, *θ*_*q*_ for *H*_2_,

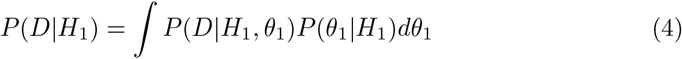

and

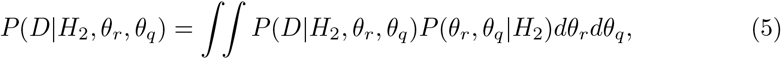

from which the Bayes factor, *BF*_12_ = *P* (*D* | *H*_1_)*/P* (*D* | *H*_2_). can be computed. This integration is typically computationally intensive, involving Monte Carlo ([23]) or related approaches such as nested sampling [24, 25]. As several tens of thousands of such numerical integrations may need to be evaluated when comparing entire transcriptomes, we use the Bayesian Information Criterion (BIC) heuristic [21, 22] as a proxy for Bayes factors. BIC is defined as

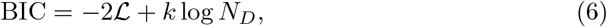

where ℒ = max_*θ*_[log *P* (*D*|*H, θ*)] is the maximised log-likelihood for *H*_1_ (model *m*_1_ with parameters *θ*_1_) or *H*_2_ (models *m*_*r*_ and *m*_*q*_ with parameters *θ*_*r*_ and *θ*_*q*_). *k* is the total number of parameters (here, 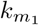 for 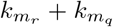 for *H*_2_), and *N*_*D*_ is the sample size (*N*_*D*_ = *N*_*r*_ + *N*_*q*_ unless some of the points overlap or a subset is selected). The lower the BIC, the better the model. Thus, if BIC(*H*_1_) *<* BIC(*H*_2_) then *H*_1_ is considered to be a better model than *H*_2_, i.e. the two time series are best explained by one model and the dynamics can thus be considered to be similar.

### 2.2 Two time series over different timescales can be transformed to compare their dynamics

If the timescales over which we wish to compare gene expression do not match, we may first need to transform one dataset to find commonalities. This ca n be achieved by identifying a transformation of time such that the similarity between the time series is the greatest. We thus aim to identify a function, *h*(*t*_*q,j*_, *β*) with parameters *β*, such that the difference between the query and reference datasets is m inimal. We u se *k*_*h*_ to denote the number of parameters of *β* in the transformation function *h*. To ensure that the order of the time points is not changed, we require that *h*(*t, β*) is a monotonic function of *t*. We denote the transformed time points by 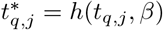.

For *H*_1_, the likelihood now evaluates the model against the transformed query dataset, 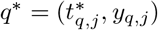,

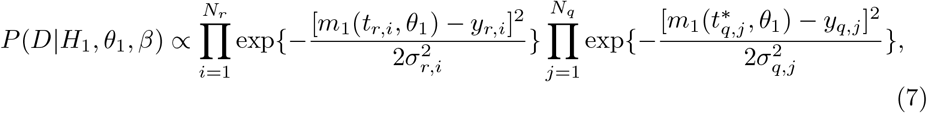

where *m*_1_(*t, θ*_1_) is the joint model of the reference and *transformed* query dataset (the transformed time points 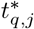 with the original expression values *y*_*q,j*_).

The additional parameters in the likelihood can be accounted for in the BIC value for 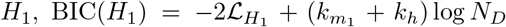, whereas BIC(*H*_2_) is unchanged, 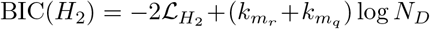, as *H*_2_ does not depend on the transformation function *h*.

To enforce monotonicity (which ensures that the order of time points remains constant) while providing a model that is simple to interpret, we chose a linear transformation with two (*k*_*h*_ = 2) biologically meaningful parameters, *β* = (*β*_1_, *β*_2_), such that *h*(*t, β*) = *β*_1_ + *β*_2_*t. β*_1_ represents a shift in time *β*_2_ and a change in timescale, i.e. a stretch factor.

Choosing a cubic spline with one knot (5 model parameters) to describe the expression dynamics, leads to 5 + 2 = 7 parameters (5 parameters for the model and 2 parameters for the transformation) for hypothesis *H*_1_ and to 5 + 5 = 10 parameters (5 parameters each for the models for reference and query) for hypothesis *H*_2_. The difference in BIC values between hypothesis *H*_1_ and *H*_2_ can be used to evaluate the similarity between the two time series. If values for the transformation parameters *β*_1_ and *β*_2_ can be found such that one joint model explains the data better than two models, BIC(*H*_1_) − BIC(*H*_2_) *<* 0, then we say the expression profiles can be *registered*. The greater the difference in BIC values, the stronger the statistical evidence. A BIC difference of 0 to 2 is considered weak evidence, 2 to 6 as positive evidence, and a difference greater than 6 as strong [22]. The maximisation of the difference in BIC values can be achieved using well-established optimisation techniques.

### 2.3 Maximisation of differences in BIC values can identify optimal transformations between two time series

To determine whether two time course datasets can be registered, we compute the difference in BIC values of *H*_1_ and *H*_2_. The BIC values depend on the model (here we use cubic B-splines with one knot based on a survey over different models for various gene expression time series) and the transformation function (here we use a linear function). The likelihood *P* (*D* | *H*_2_) and BIC(*H*_2_) do not depend on the transformation parameters and, as the number of parameters does not change, maximising the BIC difference corresponds to minimising BIC(*H*_1_). This leads to maximising the likelihood *P* (*D|H*_1_) over the model parameters *θ* and the transformation parameters *β*.

If transformation parameter values can be found that lead to the difference BIC(*H*_1_) − BIC(*H*_2_) being negative, then the data are consistent with one common model. The smaller BIC(*H*_1_) compared to BIC(*H*_2_), the more confident we can be in this inference. As explained earlier, the results depend on the choice of functions.

### 2.4 Simulated data can be used to evaluate the performance of curve registration

To create simulated datasets, we generated expression profiles using cubic B-splines for which we chose the parameters by drawing random samples from a uniform distribution *U*(−10, 10). Each resulting curve was rescaled to be positive and fall within a range of 0 to 10. For the reference dataset, ten points were chosen, *t*_*r,i*_ = 1, …, 10, from each simulated profile. For a positive control, we generated query data by sampling at ten different time points *t*_*q,j*_ = 1, …, 10 from the same cubic spline that was used to generate the reference dataset. Noise was added (from 0% to 200% using a uniform distribution) and then transformed using shift and stretch values, also sampled from a uniform distribution, shift *β*_1_ ~ *U*(0, 5) and stretch *β*_2_ ~ *U*(1*/*5, 1). For the negative control, we generated a query dataset by vertically reflecting the reference datasets around the horizontal axis to obtain curves with different dynamics to the original data. As for the positive query profiles, stretch and shift factors were applied to obtain the final negative query profiles. To evaluate the impact of the number of time points, additional sets of simulated data were generated for the positive and negative controls using five, six, seven, eight, nine, fifteen, and twenty time points. A total of 1,000 reference profiles were constructed for both positive and negative controls. These datasets allow us to evaluate our methodology over a range of conditions for which we know the correct values.

### 2.5 Implementation in R

We implemented the above approach in R. It is available as an R package *greatR* (Gene Registration from Expression and Time-courses in R). The function `register()` is used to perform the registration. Three optimisation methods, Nelder–Mead (NM) [26], L-BFGS-B [27], and Simulated Annealing (SA) [28], are available in *greatR* using the R-packages *neldermead* [29], *stats* [30], and *optimization* [31]), respectively. The default optimisation method is NM. If the parameter `optimise_registration_ parameters` is set to `FALSE`, registration will be performed using user-specified parameters, otherwise parameter optimisation is carried out by default. The output of `register()` is a list of data frames that can be used for further analysis, such as plotting using the `plot_registration_results()` function and obtaining the summary of the results using the `summarise_registration()` function. Version 2.0.0 of the package is available on GitHub at https://github.com/ruthkr/greatR/ and is distributed under the GPL-3.0 license (GNU General Public License v3.0). Users can download and install it, and learn how to use it with articles provided on https://ruthkr.github.io/greatR/.

## 3 Results

### 3.1 Validation using simulated data shows that curve registration with *greatR* correctly identifies time series with similar dynamics

To evaluate our methodology, we first generated positive control datasets using models with defined parameter values (see Methods). Our approach successfully registered all pairs with no false negatives in the noise-free case, Figure 2. We then evaluated the impact of noise in the data. To assess the contribution of noise, we first deliberately did not account for noise in the model, i.e. not adjusting the *σ* values in the likelihood for different noise levels. As expected, as the noise level increases, the registration success decreases, dropping to about 25% correct for for noise levels above 100%, Figure 2. This trend is consistent across different numbers of time points. This is expected as the dynamics become more dissimilar with increasing noise levels, favouring different models (hypothesis *H*_2_) over one model (hypothesis *H*_1_). Examples of the impact of varying noise levels are shown in Figures 3 and 4. Although many pairs could still be registered at high noise levels, the estimated stretch and shift parameters deviated from the true parameters (Supplementary Figures S2, S3, S4, S5). However, if the noise is correctly accounted for in the likelihood to match to actual noise level, i.e. the noise can be estimated from the data, we can successfully identify all positive controls Figure 2.

**Figure 2:**
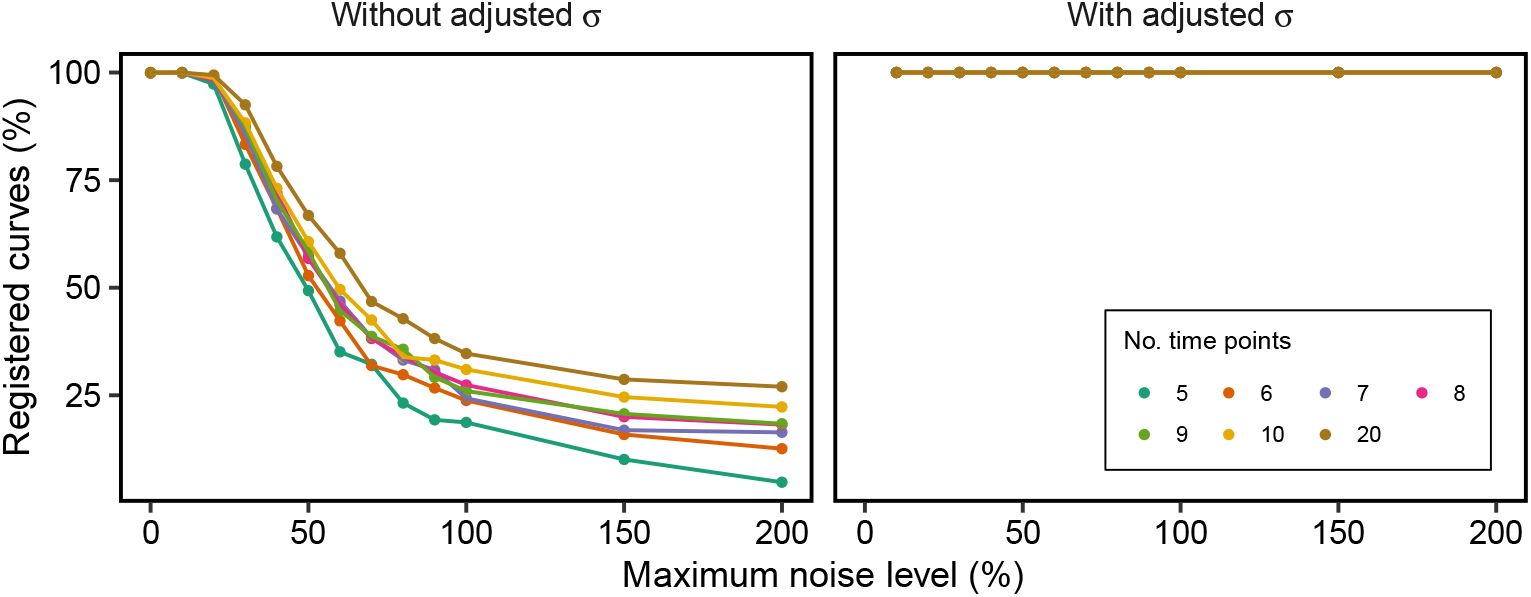
The impact of noise on curve registration. Noise can lead to a drop in performance if the noise is not taken into account in the likelihood (left plot). Total percentage of registered curves on the positive simulated data with different levels of noise and number of time points. The stronger decrease in performance for a higher number of time points is due to the more complex patterns that arise in the data. Including the estimated noise level in the likelihood function corrects this behaviour (right plot). Each point in the graph is computed over a dataset of 1,000 pairs of simulated time series.

**Figure 3:**
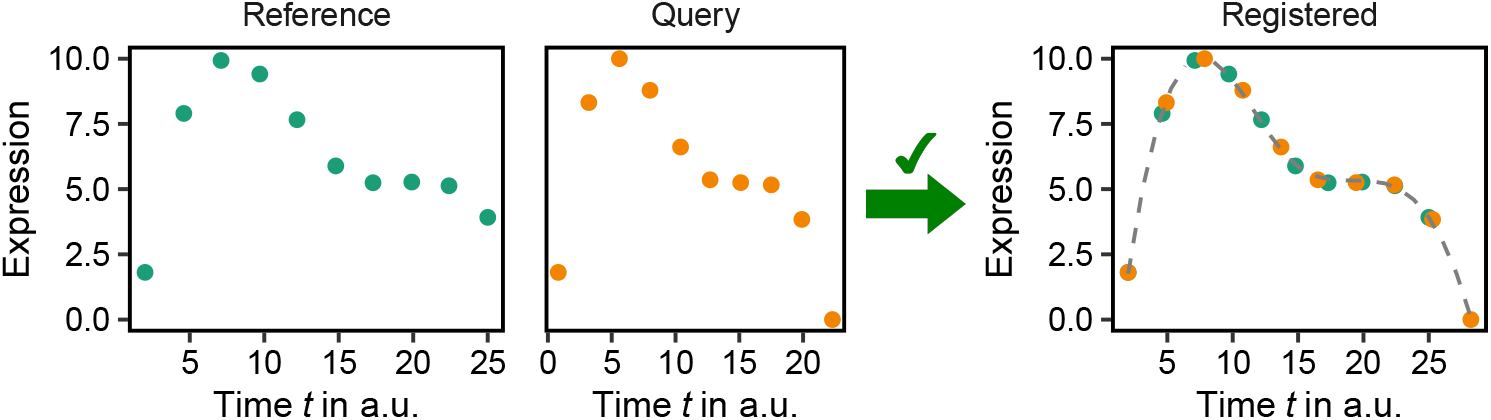
An example from the positive control dataset (with no noise). The left-hand side shows the original dynamics of both reference and query curves, and the right-hand side shows the registered curves. Green and orange indicate reference and query data, respectively.

**Figure 4:**
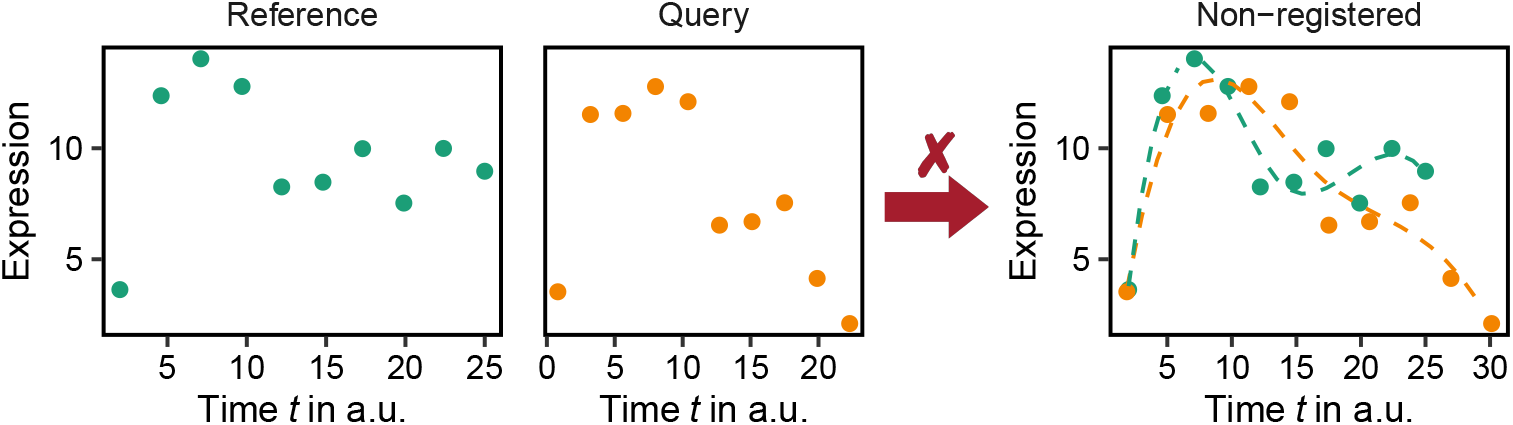
An example from the positive control dataset (noise level is equal to 60%) identified as non-registered curves. The left-hand side shows the original dynamics of both reference and query curves. The right-hand side shows the best fit of the corresponding data but in this case the curves were not deemed to be similar (not registered). Green and orange indicate reference and query data, respectively.

### 3.2 Validation using simulated data shows that curve registration with *greatR* identifies time series with different dynamics

Having established that similar dynamics can be identified using curve registration, we generated a negative control dataset to evaluate the potential of this method to find false positives. 99.6% of the negative control datasets with ten time points were correctly identified as having different dynamics. The 0.4% false positives that registered despite being from different models are due to the ability of *greatR* to identify local similarities (see Supplementary Figure S6). If we require that the transformed datasets must overlap completely in time, then we correctly identify all datasets with different dynamics with no false positives. Figure 5 shows an example of the original dynamics and the registration results of a pair of curves with different dynamics in the negative control dataset which is identified as non-registered by *greatR*. Similar behaviour was observed when the number of time points was reduced to only five. When no restrictions are applied, only 98.4% of the total number of curves were identified as having different dynamics, whereas none of them are registered when only global similarity is considered.

**Figure 5:**
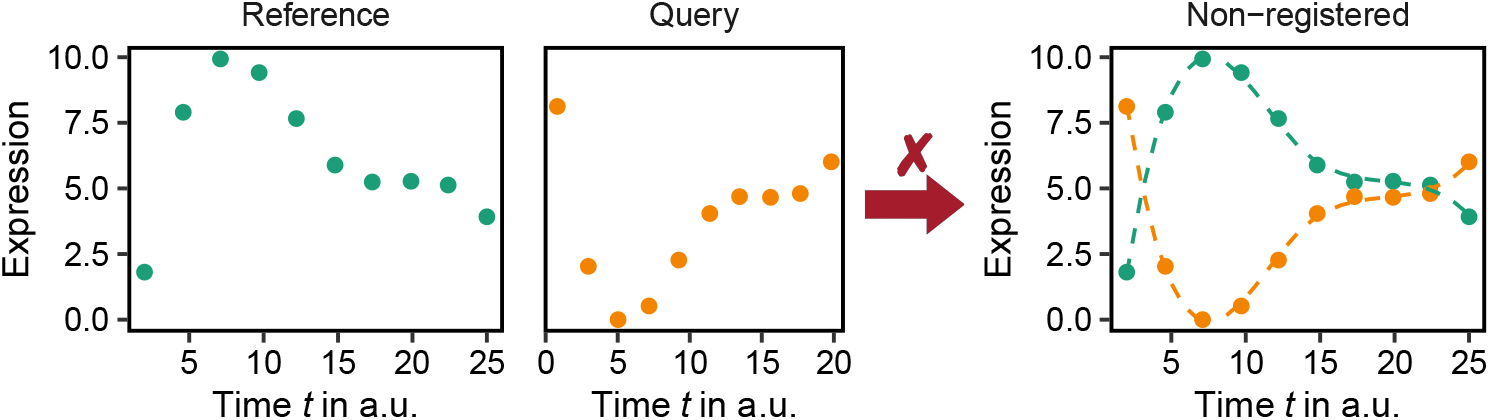
An example from the negative control dataset identified as non-registered curves. The left-hand side shows the original dynamics of both reference and query curves. The right-hand side shows the registration results of the corresponding data. Green and orange indicate reference and query data, respectively.

Using both positive and negative control datasets, we evaluated the performance of the presented methodology to identify similar time series and infer their shift and stretch factors. Our approach correctly aligns datasets from the positive control dataset and successfully identifies curves from negative control dataset as being different. We found this approach to be robust with respect to noise of up to approximately 200%, beyond which the performance deteriorates.

### 3.3 Comparative transcriptomics using *greatR* to explore expression dynamics during the floral transition between Arabidopsis, *Brassica rapa*, and *Brassica oleracea*

We now present an example using real data from a study on flowering time in Brassica crops. Among the Brassica species, *B. rapa* and *B. oleracea* are two important representative vegetable crops. Whole genome triplication played an important role in the speciation and morphotype diversification of these crops [32], resulting in genes being present in multiple copies in the genome. Gene duplication can lead to multiple outcomes: duplicated genes may accumulate deleterious mutations and become nonfunctional, or they may acquire advantageous mutations, resulting in new functionality, sub- and neofunctionalization [33]. To gain insights into the variation at the transcriptome level, we compared RNA-seq time-courses of *B. rapa*, and *B. oleracea* with the model organism *Arabidopsis thaliana*. Detecting differences in expression dynamics is important as these may potentially lead to subfunctionalization or neo-functionalization, and contribute to phenotypic differences in flowering time regulation between the three species. Here, we describe how curve registration can be exploited to compare the dynamics of genes associated with flowering time between different species. Identifying similar dynamics provides a means of transferring knowledge from the model plant *Arabidopsis thaliana* to other, lesser studied species. We used *B. rapa* cv. R-o-18 gene expression data [19], and *B. oleracea* expression data under the same growth conditions and collected from the apex over development during the vegetative growth and the floral transition, continuing until floral buds were visible (developmental stage BBCH51). For *B. rapa* and *B. oleracea*, BBCH51 was reached at 35 days and 51 days post-sowing, respectively. At each time point, three replicates were collected. For the model species, publicly available gene expression data in Arabidopsis Col-0 shoot apex from 7 to 16 days after germination grown under similar 16-hr day conditions were downloaded from NCBI SRA, project ID PRJNA268115 [34]. We first identified the corresponding orthologs of 306 genes involved in flowering time in Arabidopsis [35], revealing 487 genes in *B. rapa* and 524 genes in *B. oleracea*. The data cover an equivalent developmental period but are on different absolute time. Using *GreatR* to compare these Brassica paralogues to their homologues in Arabidopsis, we found that approximately 84% of the 524 in *B. oleracea* and 82% of the 487 in *B. rapa* flowering time-related genes were successfully registered. This result suggests that the transcriptomic differences between the two Brassica species and Arabidopsis are primarily due to temporal adjustments rather than inherently different expression profiles arising from changes in the underling gene regulatory networks.

#### 3.3.1 A comparative analysis of Arabidopsis orthologues with single-copy genes in *B. rapa* and *B. oleracea*

Despite the whole genome duplication events mentioned above, we found 72 orthologues of Arabidopsis identified in *B. rapa* and *B. oleracea* that exist as singletons. Such single-copy genes have been termed “duplication-resistant” genes [36] and could be important for the long-term survival of a polyploid lineage [37]. These singleton genes thus present an interesting test case for identifying which sets have similar dynamics in all three species and which ones have evolved species-specific dynamics. Applying *greatR* to the singleton genes found that a high proportion (71%) registered, suggesting conserved dynamics and function. Examples of such common dynamics are shown in Figure 6. Furthermore, we identified expression dynamics in this group that had common dynamics between *B. rapa* and *B. oleracea* but different from Arabidopsis, Figure 7. This analysis can be extended to investigate genes with expression dynamics that are specific to a single species and may provide clues to their phenotypic differences.

**Figure 6:**
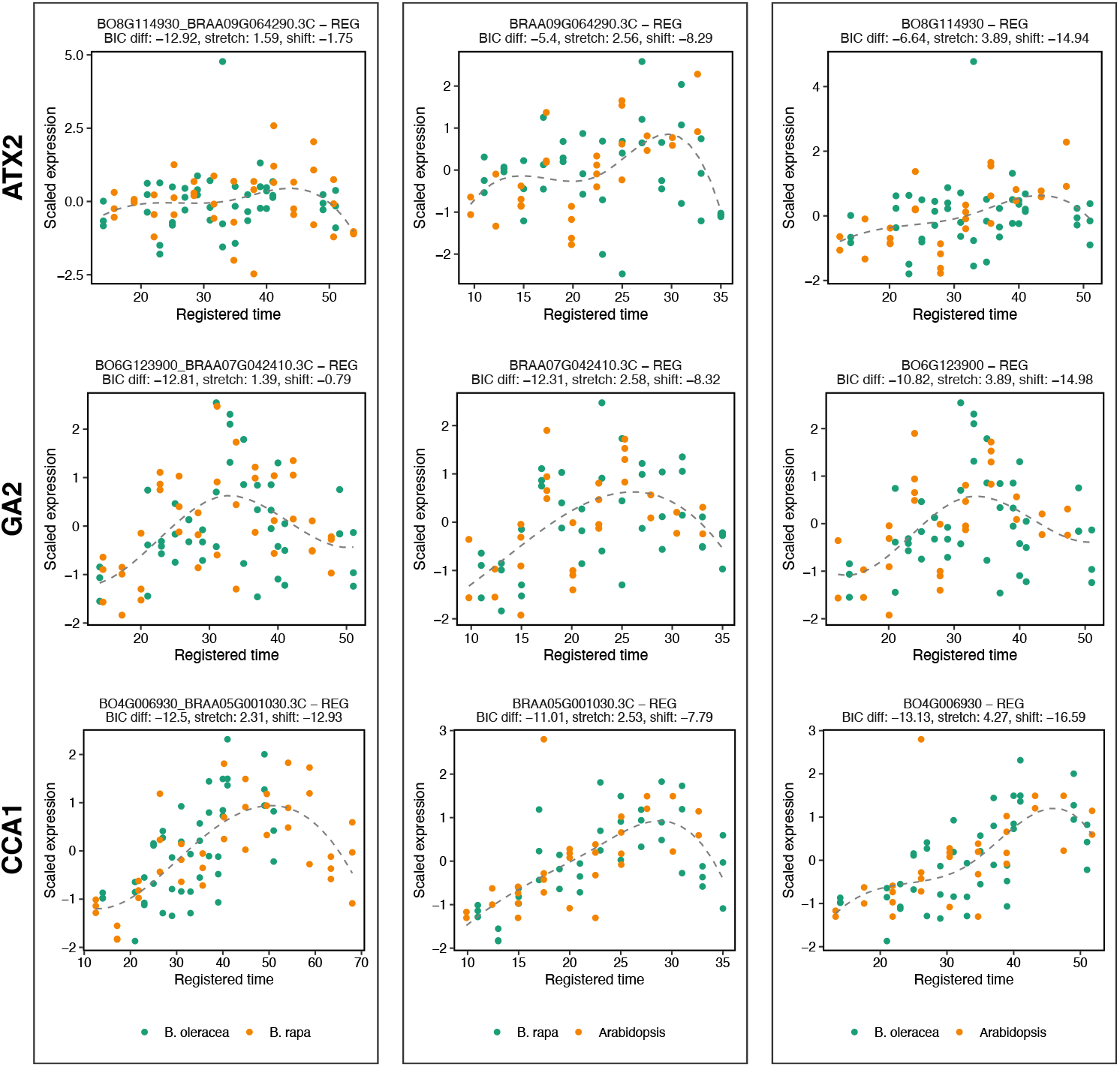
Curve registration with *greatR* can be used to compare *FLC* paralogue dynamics in *B. oleracea* and Arabidopsis. Green and orange dots represent expression replicates for each time point in *B. oleracea* and Arabidopsis, respectively. B03G005470 FLC is the only gene that did not register to its Arabidopsis homologue.

**Figure 7:**
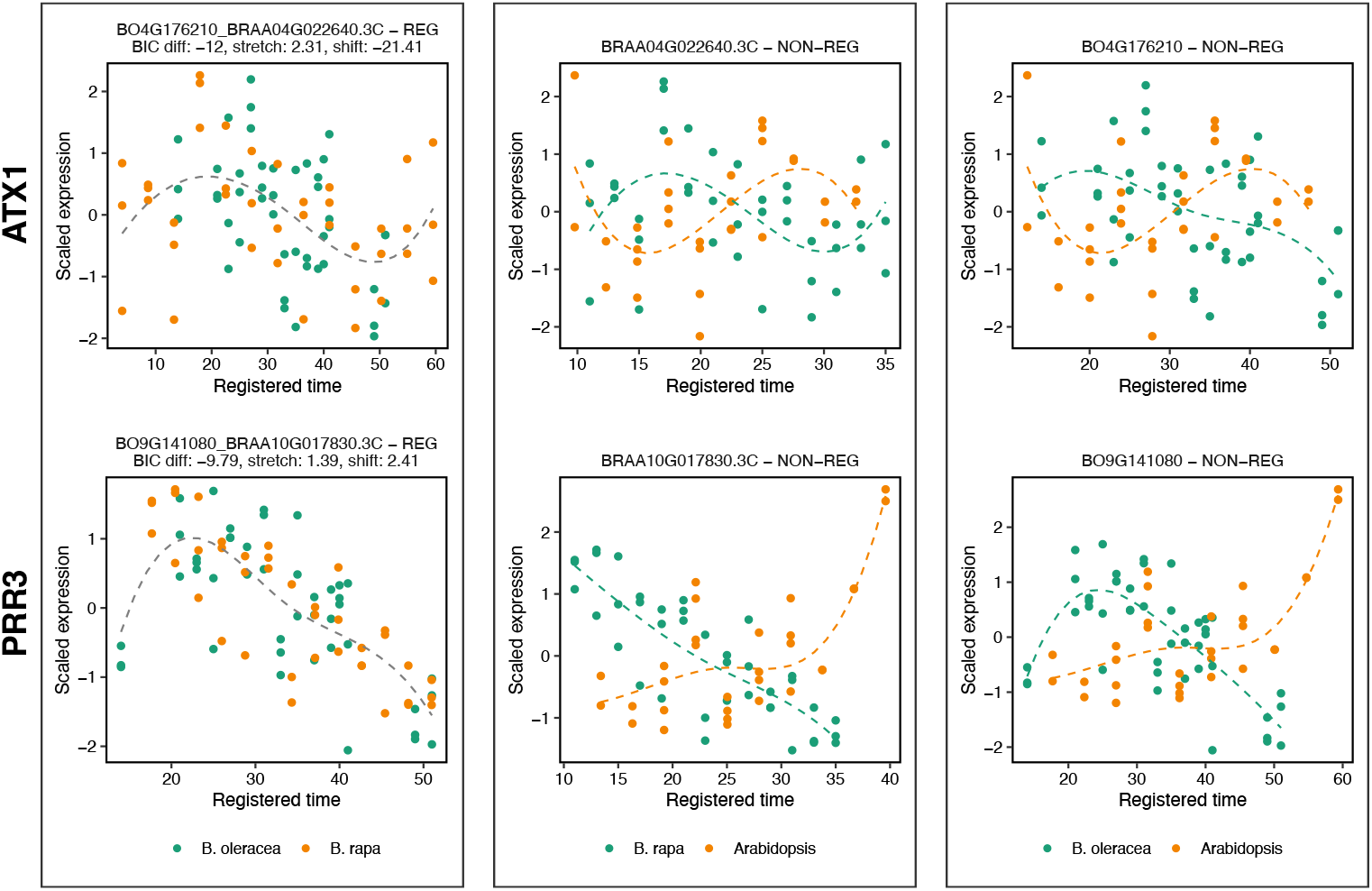
*greatR* can be used to identify lineage-specific expression dynamics. *ATX1* and *PRR3* are categorised as Brassica-specific genes. Each panel shows the pairwise comparison between species, from left to right: *B. rapa* vs. *B. oleracea, B. rapa* vs. Arabidopsis, and *B. oleracea* vs. Arabidopsis. Each dot represents a gene expression replicate at each time point. Green and orange colours denote gene expression for the species indicated in the legend below each plot. A grey dashed line indicates that the two profiles are registered as a single model fitted to both gene expression profiles. If no grey dashed line is present, the pair is not registered, with each profile fitted separately (green and orange dashed lines corresponding to the species indicated in the legend). The subtitle in each plot provides the gene ID and the optimal stretch and shift parameters for each pair of profiles.

#### 3.3.2 A comparative analysis of Arabidopsis orthologues with multiple-copy genes in *B. rapa* and *B. oleracea*

An interesting question in polyploids relates to the functionality of those genes that have been retained in multiple copies. Different expression dynamics are indicative of regulatory changes between such multiple-copy genes. We show how our approach can be employed to identify homologues that may contribute to species-specific adaptation for the flowering time gene *FLC. FLC* is a well-studied floral repressor known for its key role in the vernalization pathway. There are five *FLC* paralogues in both *B. rapa* and *B. oleracea*, so we were interested in comparing their expression profiles against the dynamics in Arabidopsis. The results of applying *greatR* to the transcriptome profiles of *FLC* paralogues are shown in Figures 8 and 9. This analysis shows how curve registration can be used to provide clues about regulatory changes and potential adaptation events by identifying which paralogue family members have similar expression dynamics to the Arabidopsis homologue and which are dissimilar.

**Figure 8:**
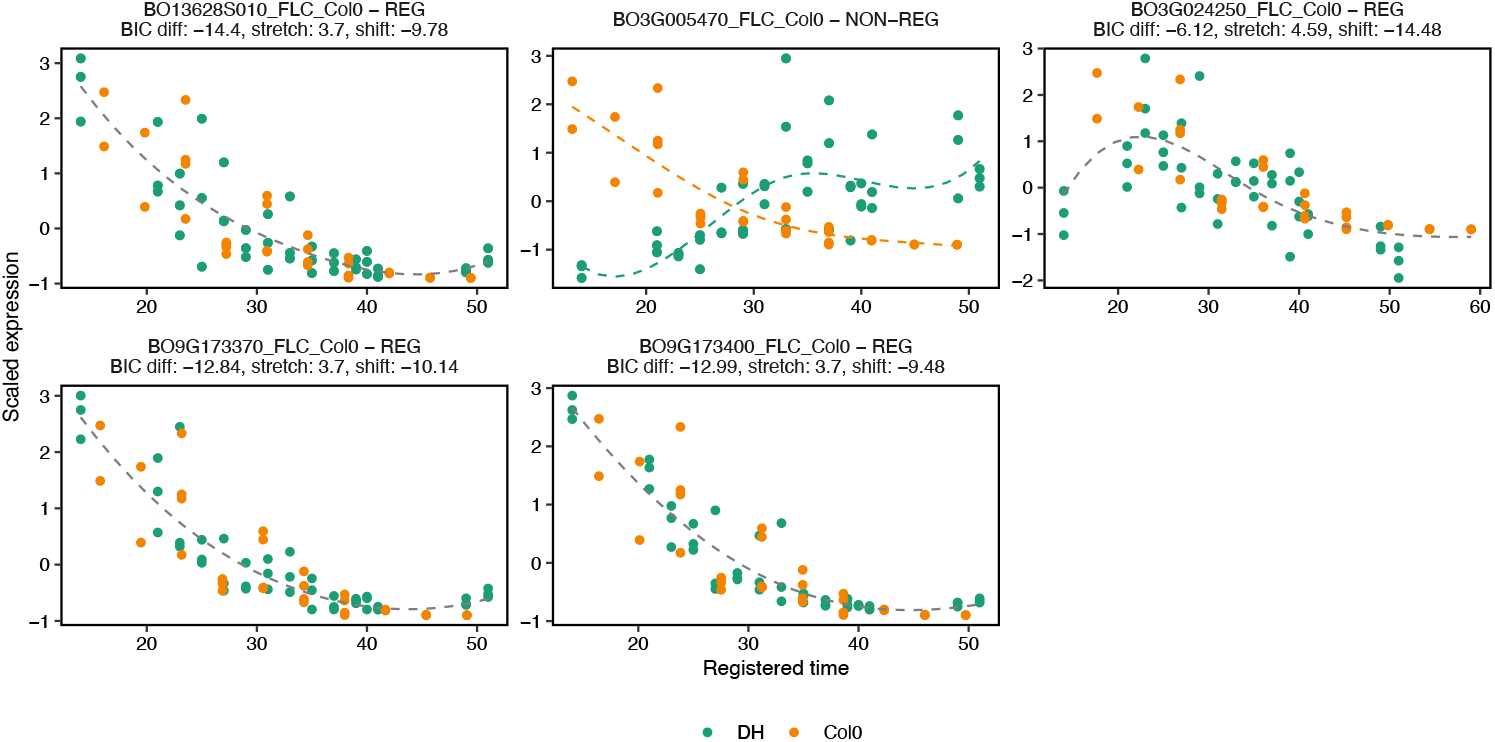
Curve registration with *greatR* can be used to compare *FLC* paralogue dynamics in *B. oleracea* and Arabidopsis. Green and orange dots represent expression replicates for each time point in *B. oleracea* and Arabidopsis, respectively. BO3G005470 *FLC* is the only gene that did not register to its Arabidopsis homologue.

**Figure 9:**
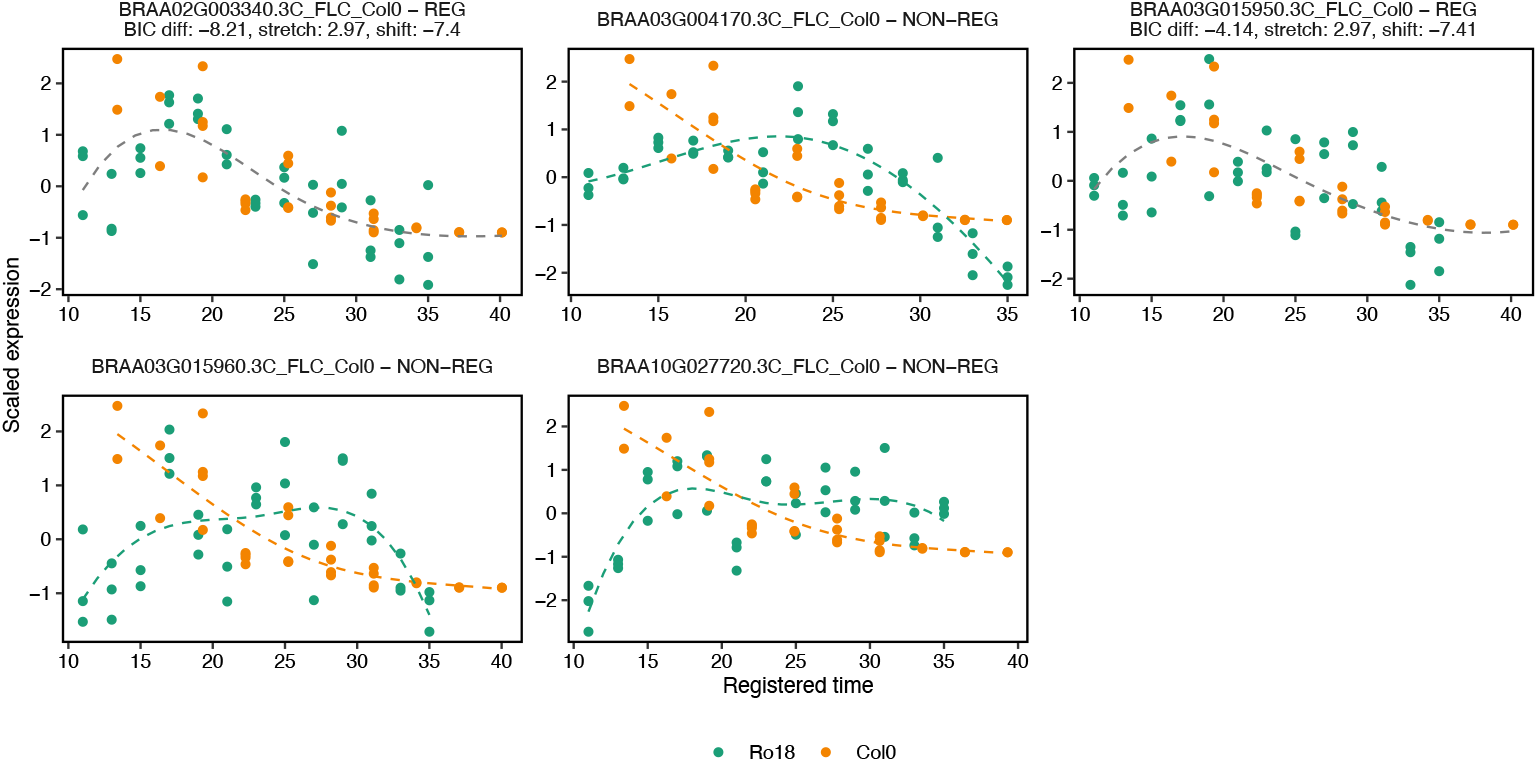
Curve registration can distinguish different dynamics between *FLC* paralogues in *B. rapa* and Arabidopsis. Out of the five identified copies of *FLC* in *B. rapa*, only two paralogues, BRAA02G003340 *FLC* and BRAA03G015950 *FLC*, were successfully registered to their Arabidopsis homologue.

## 4 Discussion

The analysis of gene expression time series holds great potential for gaining deeper insights into dynamic biological systems and processes as they unfold over time. Whilst network inference and predictive dynamic models are frequently the goal, a first question might be whether genes share similar patterns of expression or whether transcriptional responses differ under varied experimental conditions or between species. Differences in the times of sampling and different rates at which biological processes and development progress can make a direct comparison of data points challenging. To compare the dynamics of time series data requires an underlying model. This model could be a piecewise linear curve, leading to a simple interpolation scheme that allows new points to be estimated that can be compared to points in another dataset. More complex models could be polynomials of different orders, splines, Gaussian Process models, etc. or ideally a mechanistic model of the dynamical process. The selection of a suitable model is important to ensure accurate inferences. Once a model is determined, appropriate statistical analyses can be performed to assess the similarity between the pairs of datasets, now described by pairs of curves. Here, we have developed a user-friendly R package (*greatR*) that enables the comparison of different gene expression profiles based on curve registration. Our proposed approach utilises a simple linear transformation with two parameters: a stretch and a shift factor. Whilst more complex transformation functions with more parameters offer more flexibility, the parameters of a linear model have biological interpretability, with stretch accounting for differences in progression speed and shift accounting for differences in timing.

Furthermore, given the typical number of datapoints, more complex models may not be supported by the data.

Our proposed approach and implementation was validated using simulated data and applied to published datasets. The results show that *greatR* is successful in identifying whether two datasets have similar dynamics, i.e. are compatible with one generating model. The validation on simulated data showed good agreement between the estimated and true shift and stretch values. The method’s accuracy for parameter inference depends on how well the noise in the data can be estimated.

Our use of available datasets for the model plant Arabidopsis and closely related crops, *B. rapa* and *B. oleracea*, shows that conserved expression dynamics can be detected across species. However, some of the FLC orthologues do not register, suggesting regulatory changes or possibly misidentification. This highlights the potential for curve registration for studying expression dynamics in biological time course data. We described the implementation and usage of *greatR*. Future modifications and enhancements include: post-registration processes, such as clustering based on groups of shift and stretch parameters, can be applied for downstream analyses, including GO enrichment analysis; machine learning techniques, such as Gaussian Processes, may replace the current cubic B-spline fitting method to handle missing or low numbers of time points more effectively; the use of nested sampling to compute the evidence would provide an attractive alternative over the current usage of optimisation and BIC values as well as delivering estimates for the accuracy of the registration parameters. The presented methodology holds promise as a valuable tool for analysing various kinds of time course data and gaining deeper insights into differences between dynamical processes.

## 6 Acknowledgements

We thank Dr Amy Briffa (JIC) for critical reading and constructive comments on an earlier draft.

## A Supplementary Tables

**Table S1:**
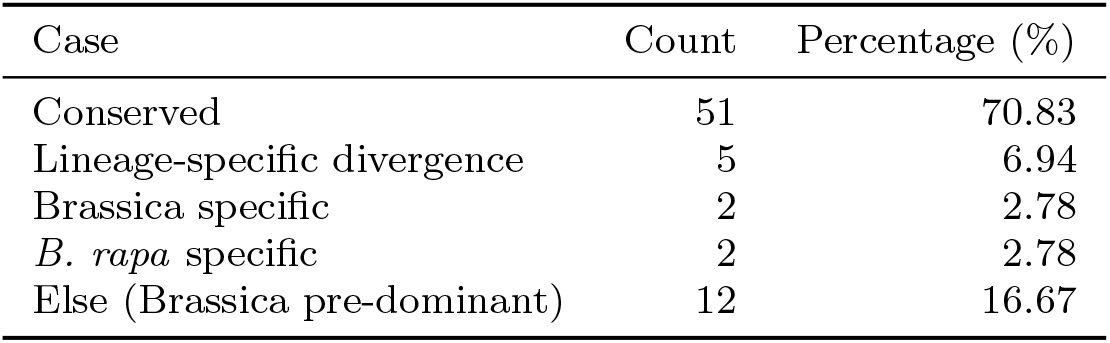
Summary of the analysis results on the conservation of single-copy gene expression through registration between three species Arabidopsis, *B. rapa*, and *B. oleracea*.

**Table S2:** BIC values for *H*_1_ and *H*_2_ across three different models, evaluated on reference and query data sets. A BIC difference of 0 to 2 is considered weak evidence, 2 to 6 as positive evidence, and a difference greater than 6 as strong [22].

| Model | BIC($H_1$) | BIC($H_2$) | BIC($H_1$) – BIC($H_2$) | Registered |
| --- | --- | --- | --- | --- |
| Linear | 2070.90 | 2046.28 | 24.61 | FALSE |
| Spline | 1152.70 | 1154.43 | -1.73 | TRUE |
| Sin | 50.07 | 55.79 | -5.72 | TRUE |

## B Supplementary Figures

### B.1 Methods

**Figure S1:**
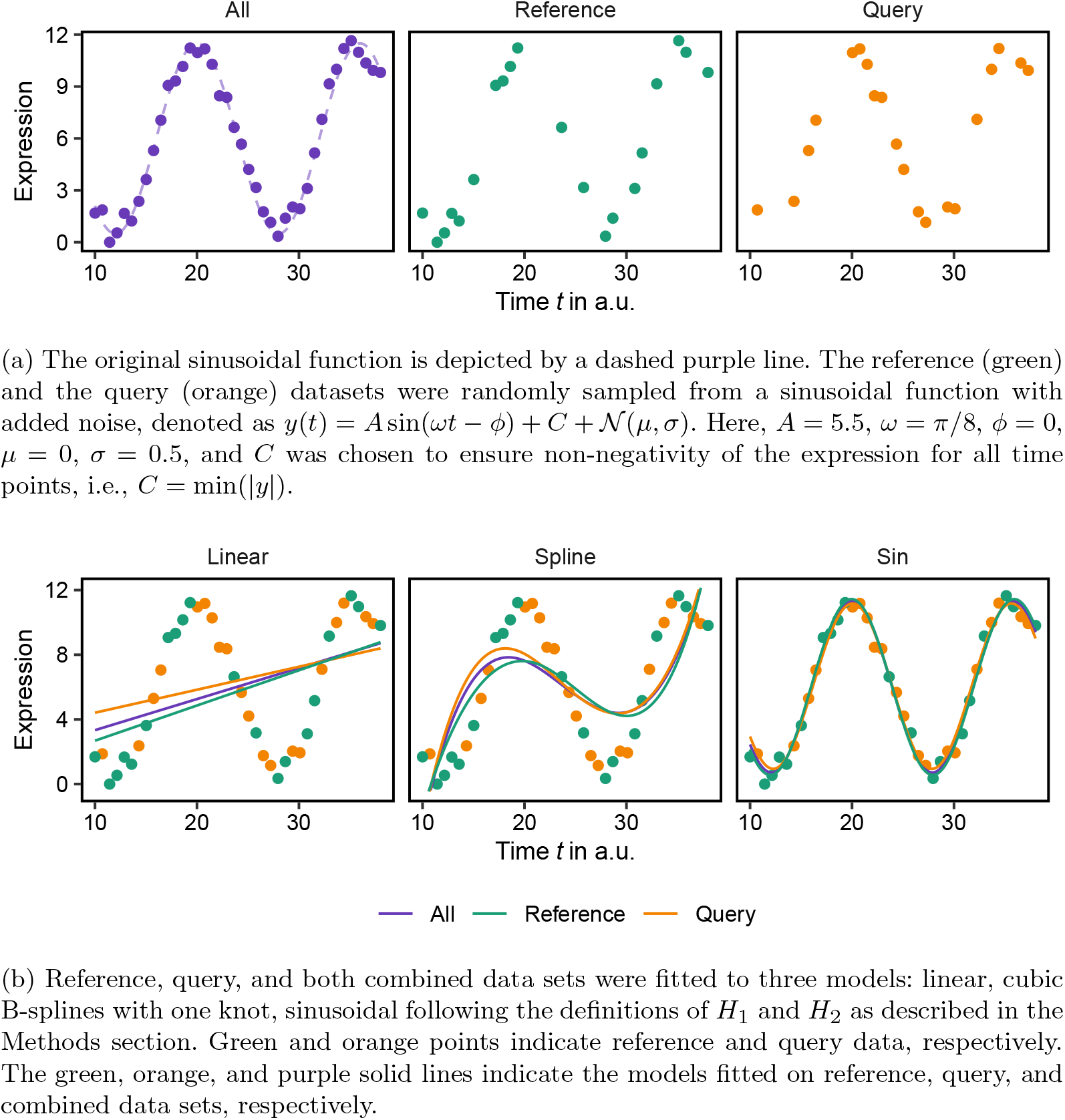
The choice of model for fitting time course data significantly impacts subsequent inferences. In the linear model, *H*_2_ is favoured over *H*_1_, indicated by the Bayesian Information Criterion (BIC) values: BIC(*H*_2_)_linear_ *<* BIC(*H*_1_)_linear_. This suggests that the query and reference data sets are better explained by two different models. In contrast, for both sinusoidal and spline models, *H*_1_ is favoured over *H*_2_, as evidenced by the BIC values: BIC(*H*_1_)_sin_ *<* BIC(*H*_2_)_sin_, and BIC(*H*_1_)_spline_ *<* BIC(*H*_2_)_*spline*_. This indicates that both reference and query data sets are best explained by a single model. See Supplementary Table S2.

### B.2 Validation using simulated data

**Figure S2:**
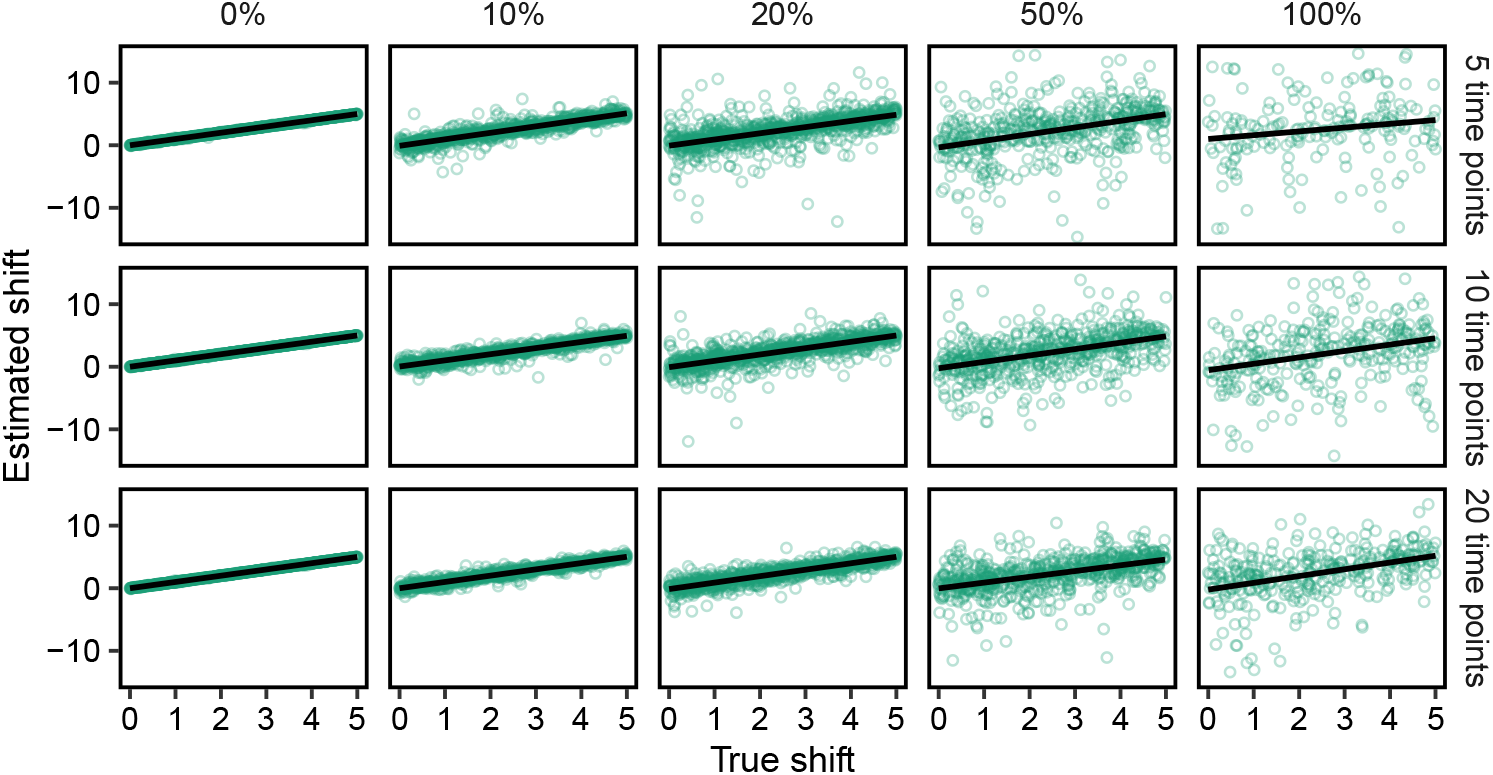
Proportion of estimated parameter shift from the registration results of simulated data based on cubic B-splines with one knot via *greatR* versus the simulated (true) value of parameter for different noise levels. Each data point represents estimation of the parameter for each gene.

**Figure S3:**
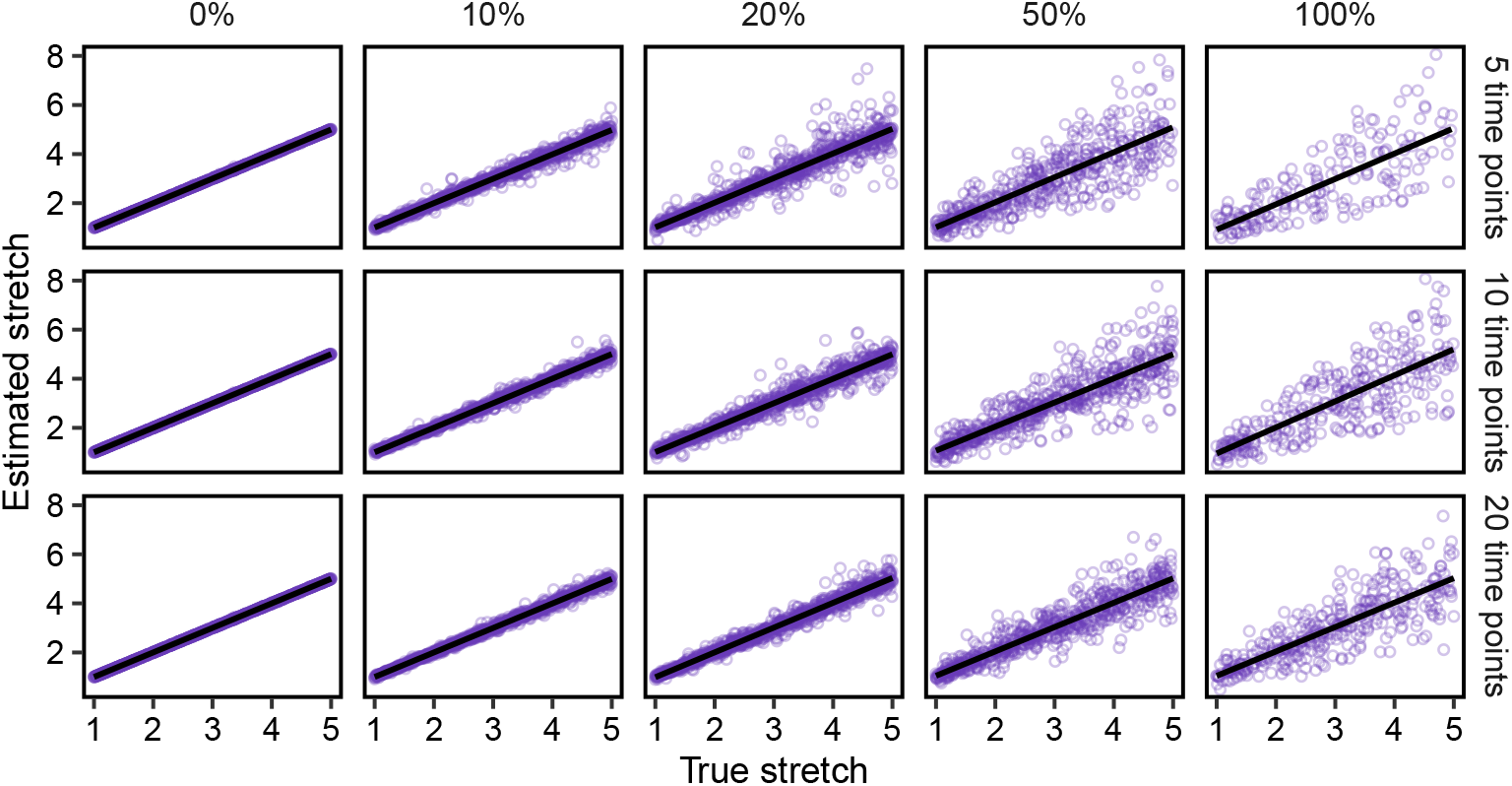
Proportion of estimated parameter stretch from the registration results of simulated data based on cubic B-splines with one knot via *greatR* versus the simulated (true) value of parameter for different noise levels. Each data point represents estimation of the parameter for each gene.

**Figure S4:**
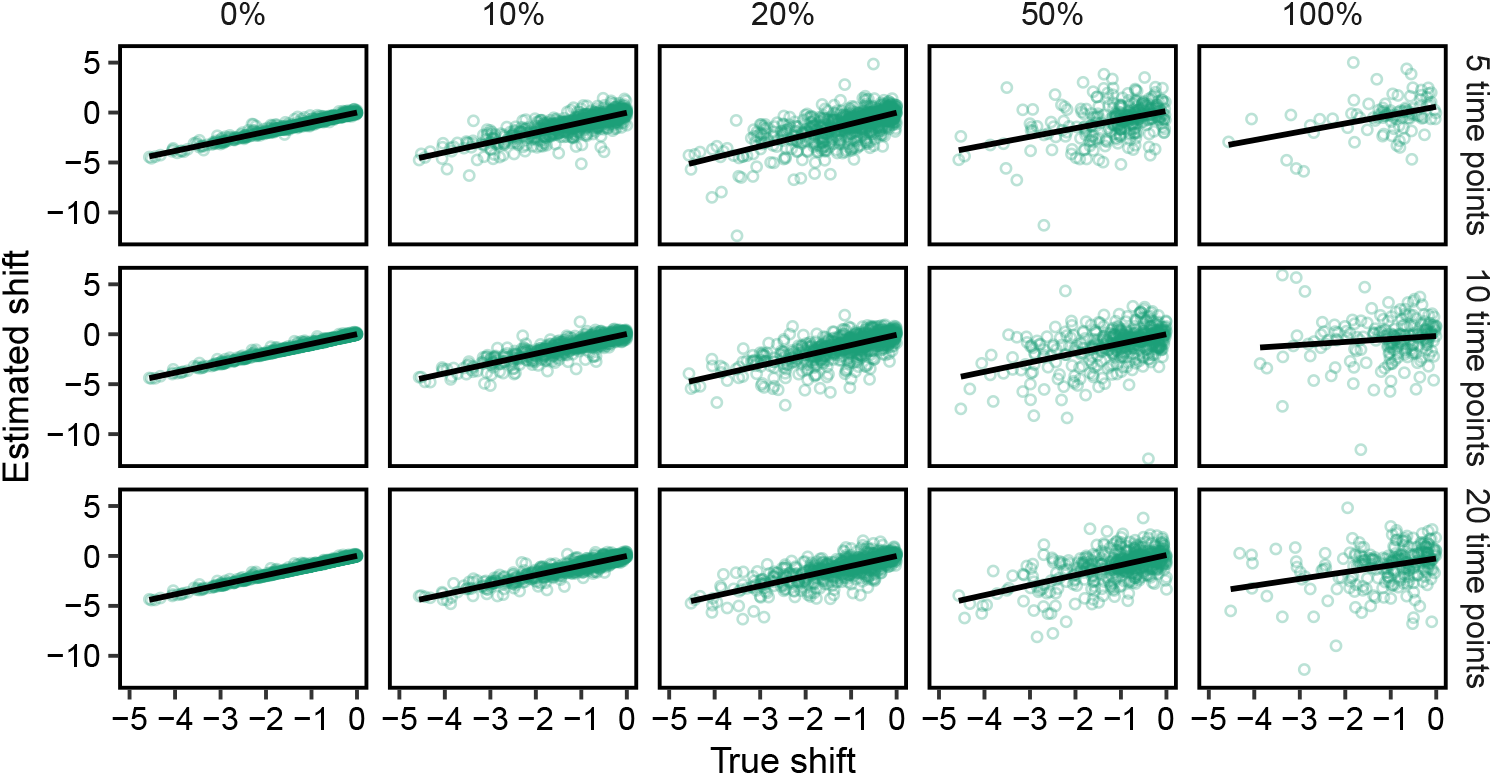
Proportion of estimated parameter shift from the registration results of simulated data based on cubic B-splines with one knot via *greatR* versus the simulated (true) value of parameter for different noise levels. The data is from the inverse transformation of Figure S2. Each data point represents an estimation of the parameter for each gene.

**Figure S5:**
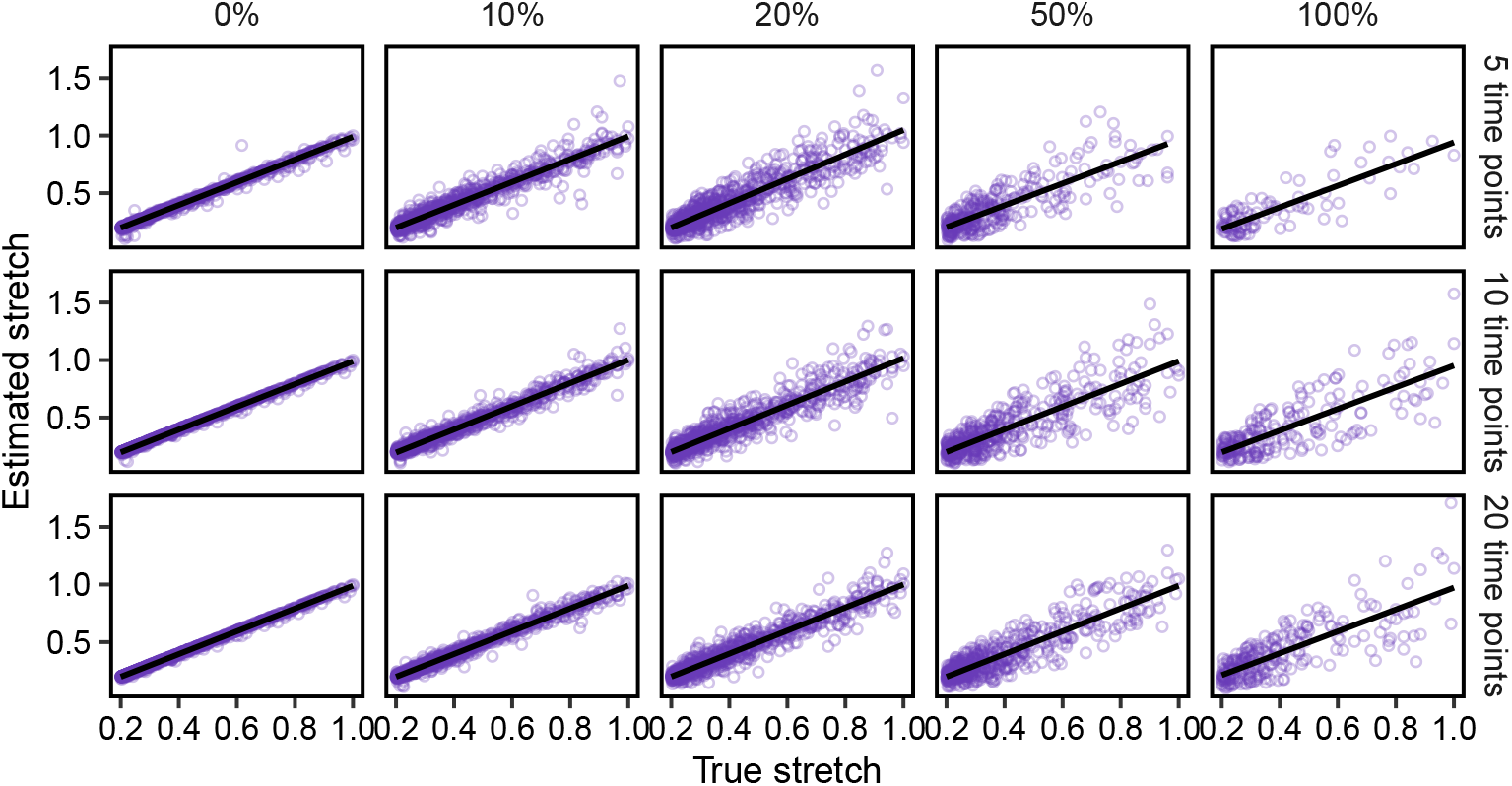
Proportion of estimated parameter stretch from the registration results of simulated data based on cubic B-splines with one knot via *greatR* versus the simulated (true) value of parameter for different noise levels. The data is from the inverse transformation of Figure S3.Each data point represents an estimation of the parameter for each gene.

**Figure S6:**
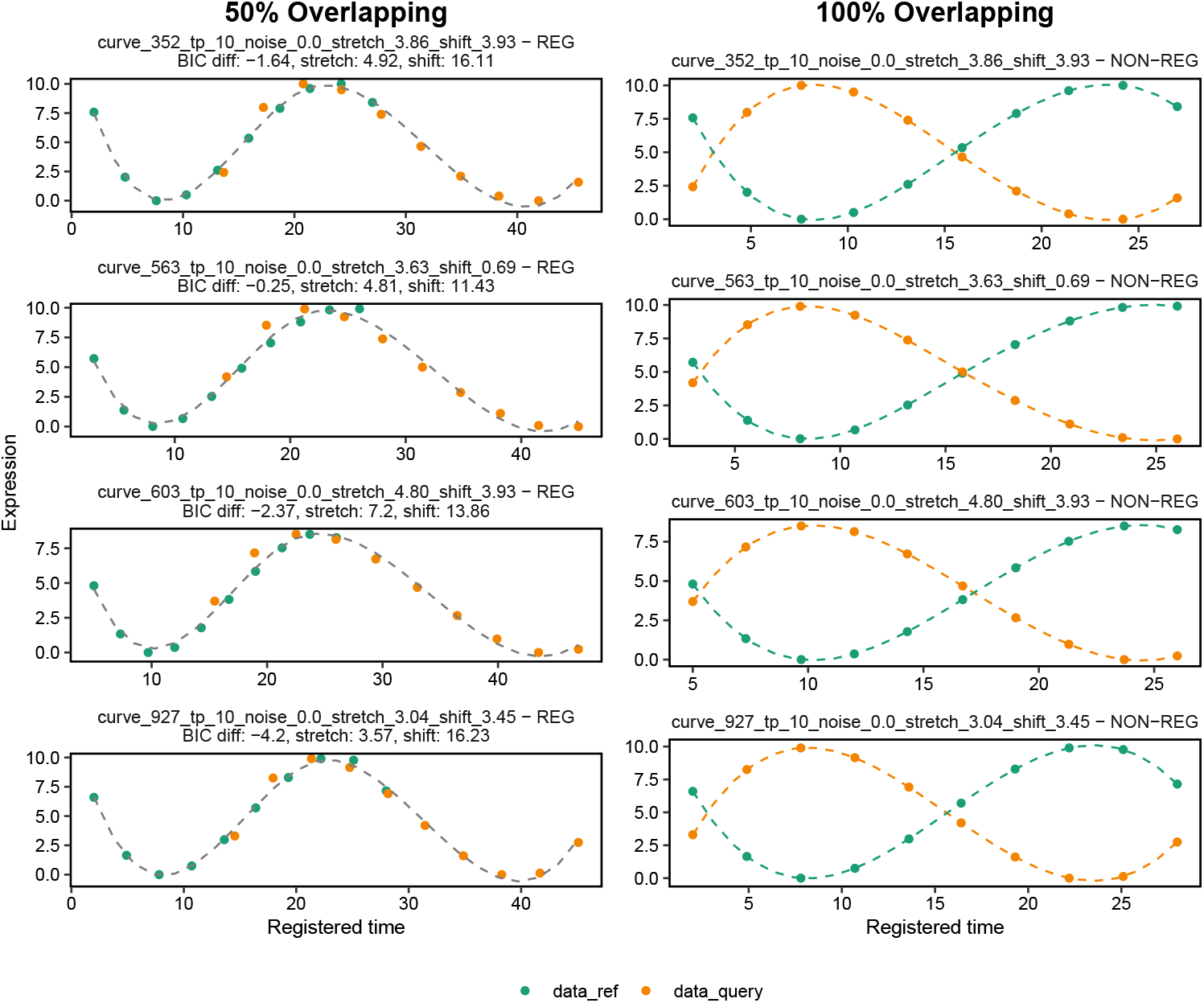
Examples from the negative control dataset where curves were initially registered due to local similarities under the 50% temporal overlap criterion. When global temporal overlap was applied, these curves were correctly identified as having different dynamics. Green and orange represent the reference and query data, respectively.

